# Ramping and phasic dopamine activity accounts for efficient cognitive resource allocation during reinforcement learning

**DOI:** 10.1101/381103

**Authors:** Minryung R. Song, Sang Wan Lee

## Abstract

Dopamine activity may transition between two patterns: phasic responses to reward-predicting cues and ramping activity arising when an agent approaches the reward. However, when and why dopamine activity transitions between these modes is not understood. We hypothesize that the transition between ramping and phasic patterns reflects resource allocation which addresses the task dimensionality problem during reinforcement learning (RL). By parsimoniously modifying a standard temporal difference (TD) learning model to accommodate a mixed presentation of both experimental and environmental stimuli, we simulated dopamine transitions and compared it with experimental data from four different studies. The results suggested that dopamine transitions from ramping to phasic patterns as the agent narrows down candidate stimuli for the task; the opposite occurs when the agent needs to re-learn candidate stimuli due to a value change. These results lend insight into how dopamine deals with the tradeoff between cognitive resource and task dimensionality during RL.

## Introduction

Since cognitive and motor resources are limited, efficient resource allocation is as important as reward maximization for animals to survive. To use their resources efficiently, animals need to focus resources on the stimuli that are highly informative of reward. However, it is not easy to know which and how many stimuli in the environment are relevant to a given task. As shown in the pigeon superstition experiment by Skinner, animals can mistakenly associate reward with a stimulus that does not predict the reward [1]. Although experimentally inserted cues (e.g. tones, light) are usually more salient and informative of a reward than other stimuli in the environment (e.g. wells, floor), pseudo-conditioning has shown that animals also link the latter with the reward [2–4]. Effective task dimensions, the task dimensions to which an animal allocates resources by approaching, learning, exploiting, exploring, or paying attention to them, often differ from the essential task dimensions (Figure 1AB). Moreover, effective task dimensions may change as learning proceeds [5–7]. Efficient resource allocation would facilitate a reduction in the effective task dimensionality during RL.

**Figure 1.**
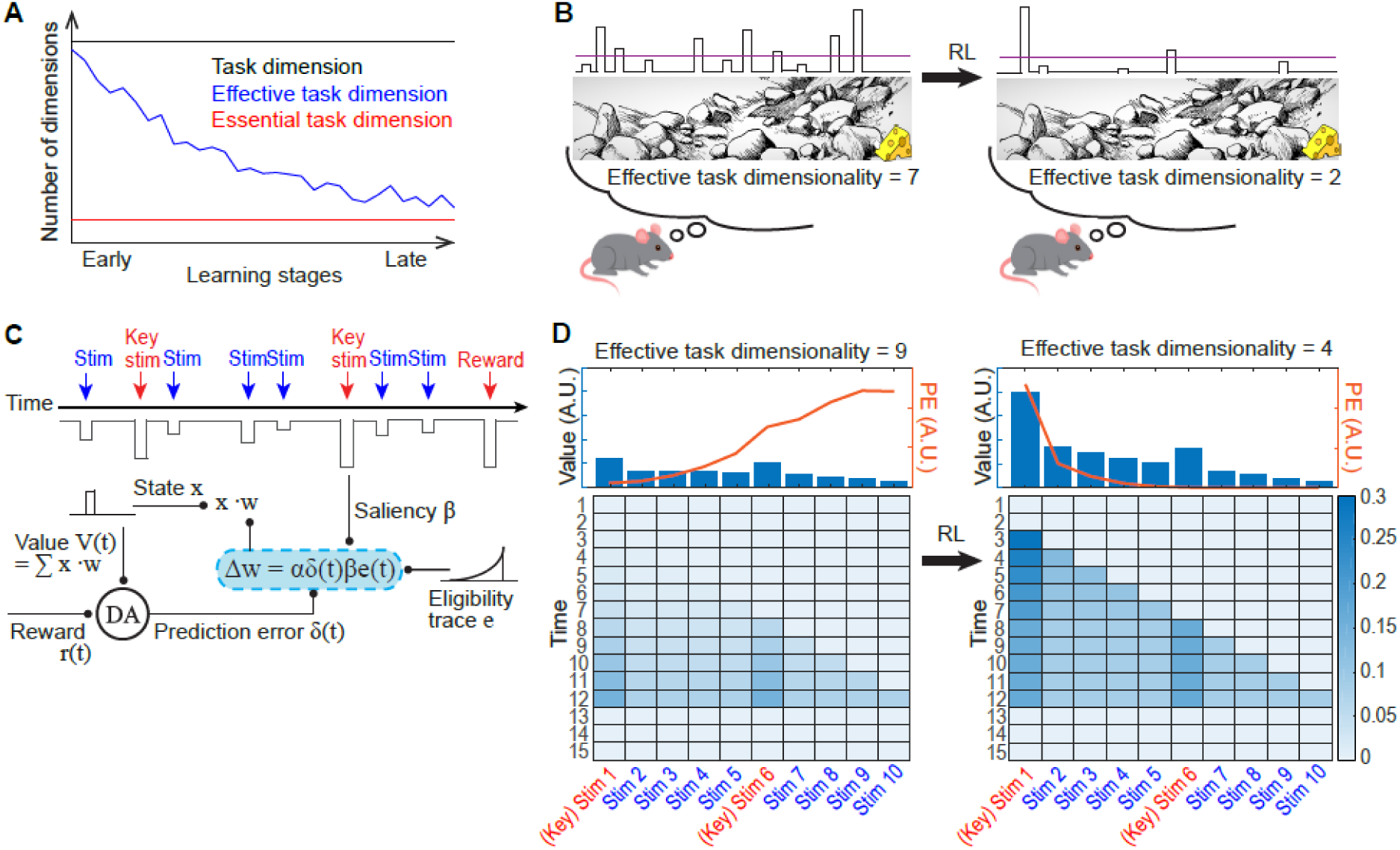
Resource allocating dopamine optimizes effective task dimensionality. (A) As learning proceeds, effective task dimensionality (the number of task variables affecting the behavior of the learning agent) is reduced to the minimum task dimensionality (the number of essential task dimensions). Task dimensions do not change in a fixed environment. (B) An illustration showing that effective task dimensionality decreases during RL in a stable environment. Animals do not perceive every stimulus in an environment, but they tend to recognize stimuli whose saliency exceed a threshold level (the purple horizontal line). (C) A modified TD model. Experimentally inserted cues (key stimuli) are usually more salient than environmental stimuli. (D) Value distribution and effective task dimensionality during early (left) and late (right) stages of learning. The bottom plot shows the weights of each stimulus over time. The top plot shows the prediction error and the weight of each stimulus summed across time. The number of stimuli which generate prediction error > 0.01 was considered as effective task dimensionality. Note that the main results still hold for other threshold values, such as 0.03 or 0.05.

Cognitive and motor resource allocation is tightly linked with an animal’s approaching, avoiding, learning, exploiting, exploring, and attending; dopamine affects all of these behaviors. In the striatum, dopamine inhibition suppresses the initiation and maintenance of instrumental responses, while dopamine excitation increases the likelihood of movement initiation [8–11]. Striatal dopamine depletion reduces high-effort, high-reward choices, while increasing dopamine has the opposite effect [12–14]. Dopamine promotes avoidance responses to aversive stimuli by modulating the activity of the brainstem-projecting prefrontal neurons and nucleus accumbens neurons [15,16]. Dopamine drives both fear learning and reward learning by signaling prediction error [17–25]. Enhancing striatal dopamine increases the exploration of novel choices while decreasing the exploitation of learned options [26–28]. Prefrontal dopamine is involved in working memory maintenance and has been suggested to reflect cognitive effort [29–31]. Dopamine neurons are necessary for the attention signal in the amygdala [32,33].

Based on the aforementioned evidence on the involvement of dopamine in resource allocation, we assumed that dopamine activity patterns are suited to managing effective task dimensionality by efficiently allocating resources to stimuli; furthermore, we assumed that the stimuli to which resources should be assigned are primarily determined by the prediction error generated by each stimulus. The role of the dopaminergic prediction error signal in RL has been much studied both experimentally and computationally. However, investigating the role of dopamine in adjusting effective task dimensionality during RL has received scant attention, partly because most RL models have been configured to learn conditioned and unconditioned stimuli only, without accommodating the effect of other environmental stimuli on task performance. Motivated by previous findings that dopamine activity may transition between ramping to phasic [34], we hypothesized that the dopamine transition from ramping to phasic supports narrowing down candidate stimuli (reducing effective task dimensionality), whereas the opposite transition helps re-learning candidate stimuli (increasing effective task dimensionality) (Figure 1AB).

To test this hypothesis, we exploited a standard class of RL models that provide predictions of stimulus values and associated prediction errors across space and time. The prediction error signals in these models have well-simulated phasic dopamine responses to experimentally inserted cues and rewards [35–38]. RL models have also provided an explanation for the gradually increasing dopamine activity which occurs as animals approach a reward, by considering internal spatial representation, the temporal decay of dopamine-dependent synaptic potentiation, the uncertainty of action timing or discounted vigor or by assuming dopamine as a value signal [39–42]. However, no RL model has accounted for the conditions in which dopamine activity transitions between ramping and phasic. To fully accommodate a mixed presentation of both experimental and environmental stimuli, we parsimoniously modified a standard TD model and simulated dopamine activity transitions between ramping and phasic (Figure 1C). By comparing the simulation results with experimental data from previous studies, we demonstrated that the dopamine transition supports the adjustment of effective task dimensionality during RL.

## Results

### The pattern of prediction error transitions from ramping to phasic as learning proceeds, reducing effective task dimensionality

Since dopamine has a resource allocating effect in its target regions, we assumed that the stimuli that evoke sufficient dopamine excitation constitute effective task dimensions. In our model, the prediction error signal simulates dopamine activity and the number of stimuli that generate a prediction error signal larger than a threshold corresponds to effective task dimensionality. To test if effective task dimensionality decreases as learning proceeds, we considered a situation in which both environmental stimuli and experimental stimuli (key stimuli in Figure 1C) were present. In this situation, frequent exposure to stimuli would shorten the effective time window during which previously experienced stimuli affected learning. To reflect this effect, a medium λ of 0.5 was used during the simulation [43–45].

The simulation shows how the shape of the prediction error signal changes over time. During the early stages of learning, values and prediction errors were widely distributed over different stimuli (Figure 1D left). In this condition, the effective task dimensionality was high and the shape of prediction error resembled the ramping pattern of dopamine activity. As learning continues, experimental cues gained substantially higher values than other environmental stimuli (Figure 1D right). The prediction error signal gradually propagated backward and was concentrated on the initial experimental cue, exhibiting a pattern similar to the phasic dopamine activity (Figures 1D and 2A). As the shape of the prediction error transitioned from ramping to phasic, effective task dimensionality decreased (Figures 1D and 2B). The simulation results suggest that during the early stages of learning, when the learning agent usually finds it difficult to identify the key stimuli, the agent temporarily exhibits ramping dopamine activity; this wide distribution of dopamine activity favors broad distributions of cognitive resources over multiple stimuli. As the agent gradually learns to identify reward-predicting stimuli, however, dopamine activity transitions from ramping to phasic, indicating that the agent focuses its resources on learning about those key stimuli.

### Limited resources and environmental stimuli account for the smooth, ramping shape of the prediction error

Previous studies have shown that even in a simple RL task, limited working memory capacity influences model-free learning [43–45,47,48]. To better understand the model behavior, we ran simulations while systematically changing λ. A large λ accommodates a situation where effective task dimensionality is smaller than the working memory capacity; this occurs when the agent has a large working memory capacity or the environment is simple enough to identify task-relevant stimuli. In this scenario, the learning agent would be able to quickly pinpoint the key stimuli. Corroborating this view, for a large λ, effective task dimensionality rapidly decreases (Figure 2B); as a result, the prediction error is concentrated on the key stimuli, which produces a phasic pattern beginning in the early stages of learning (Figure 2A). This helps the agent quickly complete learning (Figure 2B).

**Figure 2.**
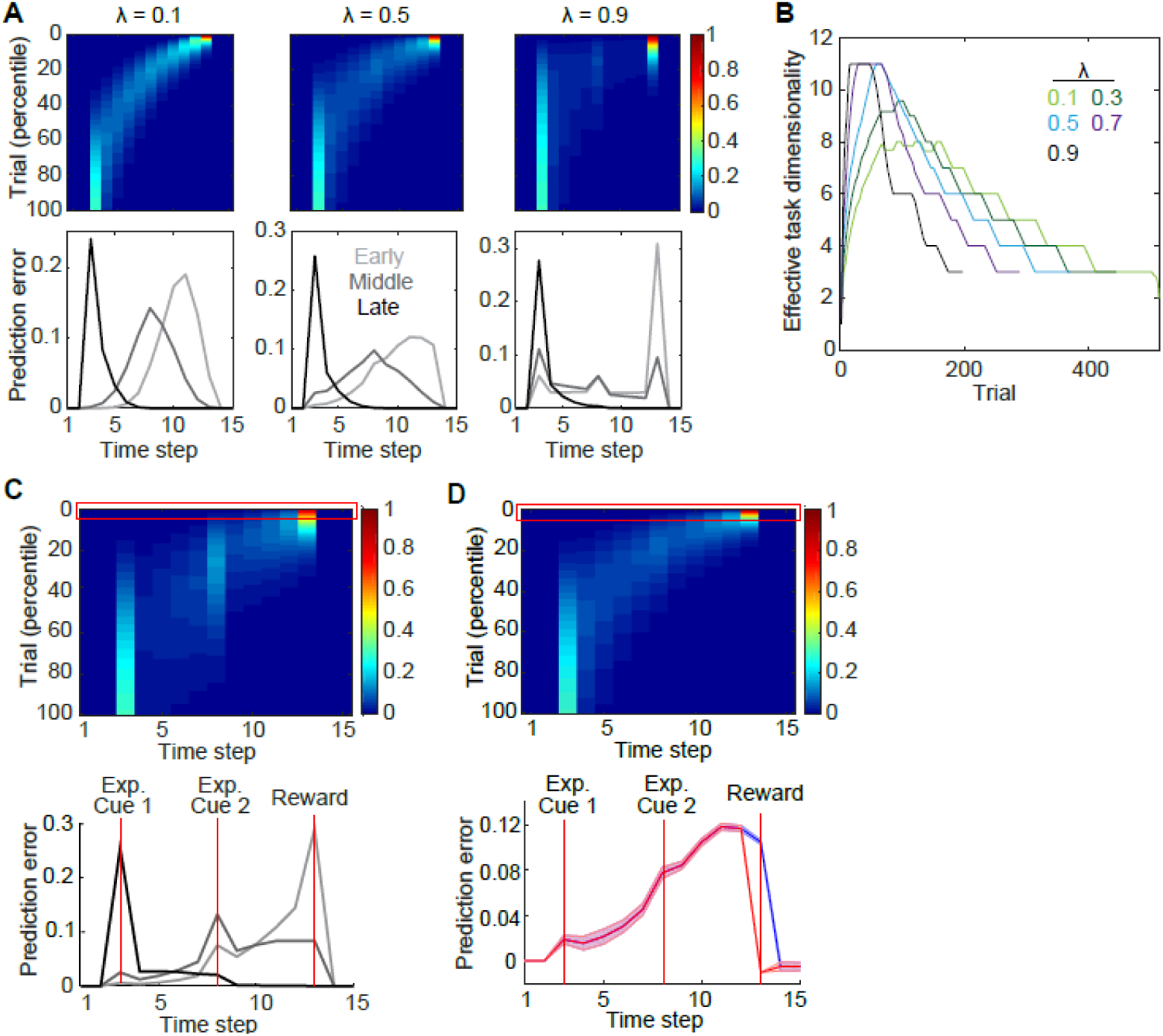
The ramping prediction error is caused by both the limit of cognitive resources and the environmental stimuli. (A) Prediction error signal during RL when λ is 0.1, 0.5, or 0.9. (B) Effective task dimensionality during RL. Stimuli that generated a prediction error larger than 0.01 were considered to constitute effective task dimension. Data until the agent completed learning is shown. The data were smoothed using a moving average window of 20. (C) The prediction error signal during RL when environmental stimuli were removed from the model. λ was 0.5. (D) The prediction error signal throughout learning (top) or during the early stage of learning (bottom) when the environmental stimuli were included in the model. Dopamine concentration in the ventral striatum during rewarded (blue) or unrewarded (red) trials. We ran 100 simulations with different initial weights (random weights with a mean of 0.02). λ of 0.5 was used. Shading shows mean ± SD.

A medium or small λ accommodates a situation in which cognitive resources can be assigned to only a portion of effective task dimensions. Reducing λ decreases the maximum size of effective task dimensionality (Figure 2B) and the width of the prediction error signal (compare the left and middle plots in Figure 2A). A wide and ramping prediction error occurs when λ has an intermediate value. The simulation results suggest that broad distributions of dopamine activity, such as ramping dopamine activity patterns, occur when effective task dimensionality is moderately smaller than the working memory capacity.

Even after the environmental stimuli are removed from the model, the prediction error shows a ramping trend if λ has an intermediate value (Figure 2C). This raises the question of whether the prediction error signal ramps up regardless of the existence of environmental cues. To answer this question, we compared our simulation results with a ramping dopamine activity reported in Howe et al. [46] (see Figure 1CE of [46]). In this study, rats were trained to travel through a large T-maze to earn a reward. The first and the second tone cues indicated the start of each trial and which arm to visit to receive the reward, respectively. A medium λ is suited to simulating this experiment because multiple environmental stimuli—such as approaching the corner of the T-maze—can provide subsidiary information to guide the animal’s behavior in a large maze, increasing the effective task dimensionality. The fact that the task performance of the animals was not very high and did not reach an asymptote suggests that the data of Howe et al. [46] may have been obtained during the early stages of learning (see Figure 4E of [46]).

When λ was 0.5, both models, with and without environmental stimuli, exhibited ramping prediction error patterns during the early stage of training. While the model with environmental stimuli replicated the observed dopamine activity (Figure 2D), the model without environmental stimuli contradicted the observed dopamine activity because the prediction error inevitably peaked at the intermediate experimental cue (Figure 2C). This finding supports our hypothesis that, when the learning agent finds it difficult to identify task-relevant stimuli, dopamine ramps up, broadly distributing resources to many candidate stimuli including experimental and environmental cues.

### Extended training transforms dopamine activity from ramping into phasic

To further validate our claim, we compared the ramping-to-phasic prediction error transition from our simulation with previous experimental results. Unlike Howe et al. [46], Collins et al. [34] trained rats for two more days after their performance reached an asymptote. In their study, the animals were trained to press two different levers to collect a reward. The authors observed that dopamine activity ramped up during early training but peaked at the first cue after extended training (Figure 3AB left). This finding is consistent with our simulation result (Figure 3AB right).

**Figure 3.**
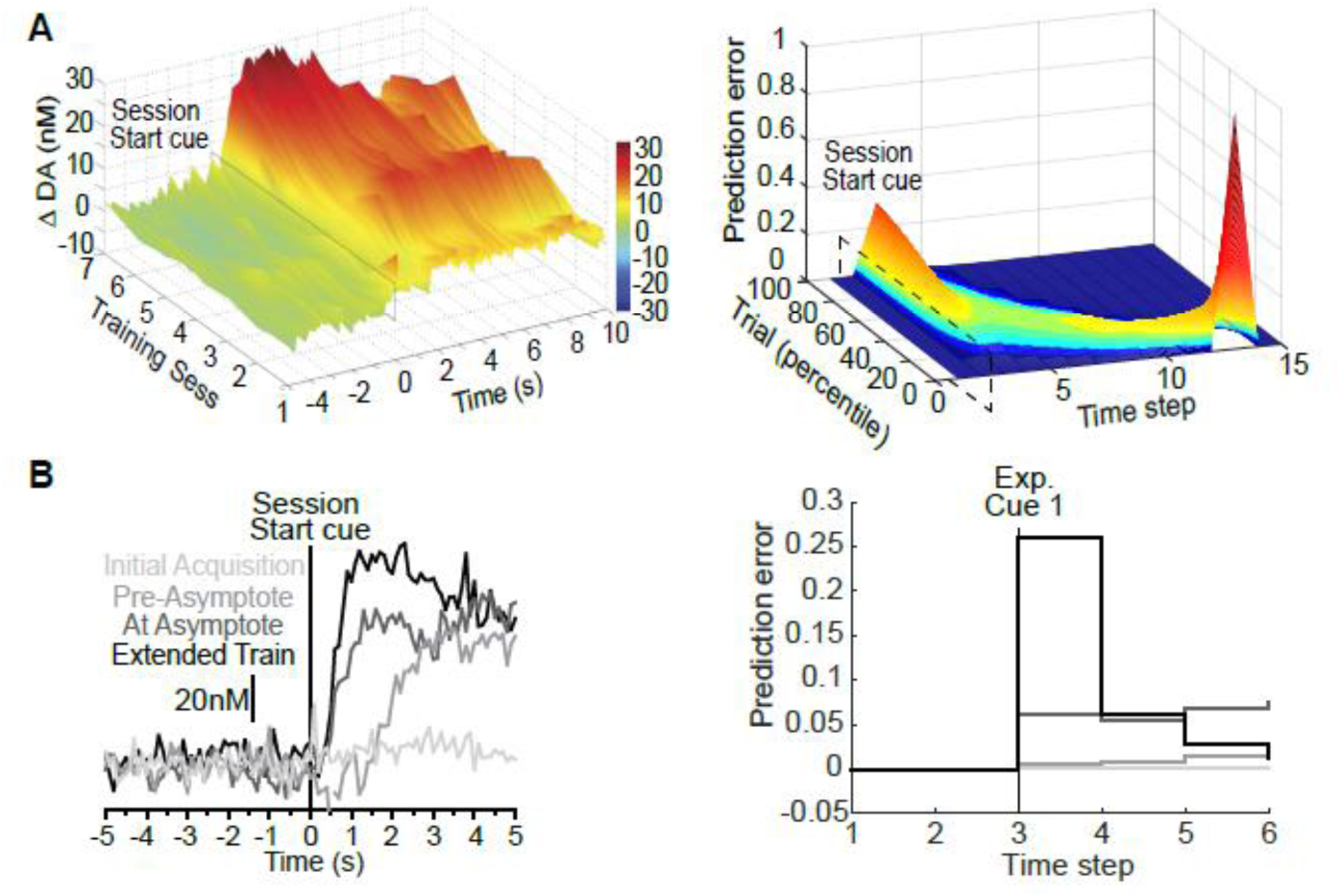
Dopamine transitions from ramping to phasic as learning proceeds. The left plots show the dopamine concentration in the ventral striatum observed in Collins et al. [34]. The right plots show the prediction error signal of the model. In (B), the 1st, 10th, 30th and 80th percentiles were considered as the initial acquisition, pre-asymptote, at asymptote, and extended training, respectively. The 30th percentile was chosen as the asymptote because the prediction error of the model peaks at the first cue around the 30th percentile. The parameter values were the same as in Figure 2E. Plots on the left were originally published in Collins et al. (2016) [34] under a CC-BY 4.0 license and are adapted here with permission.

**Figure 4.**
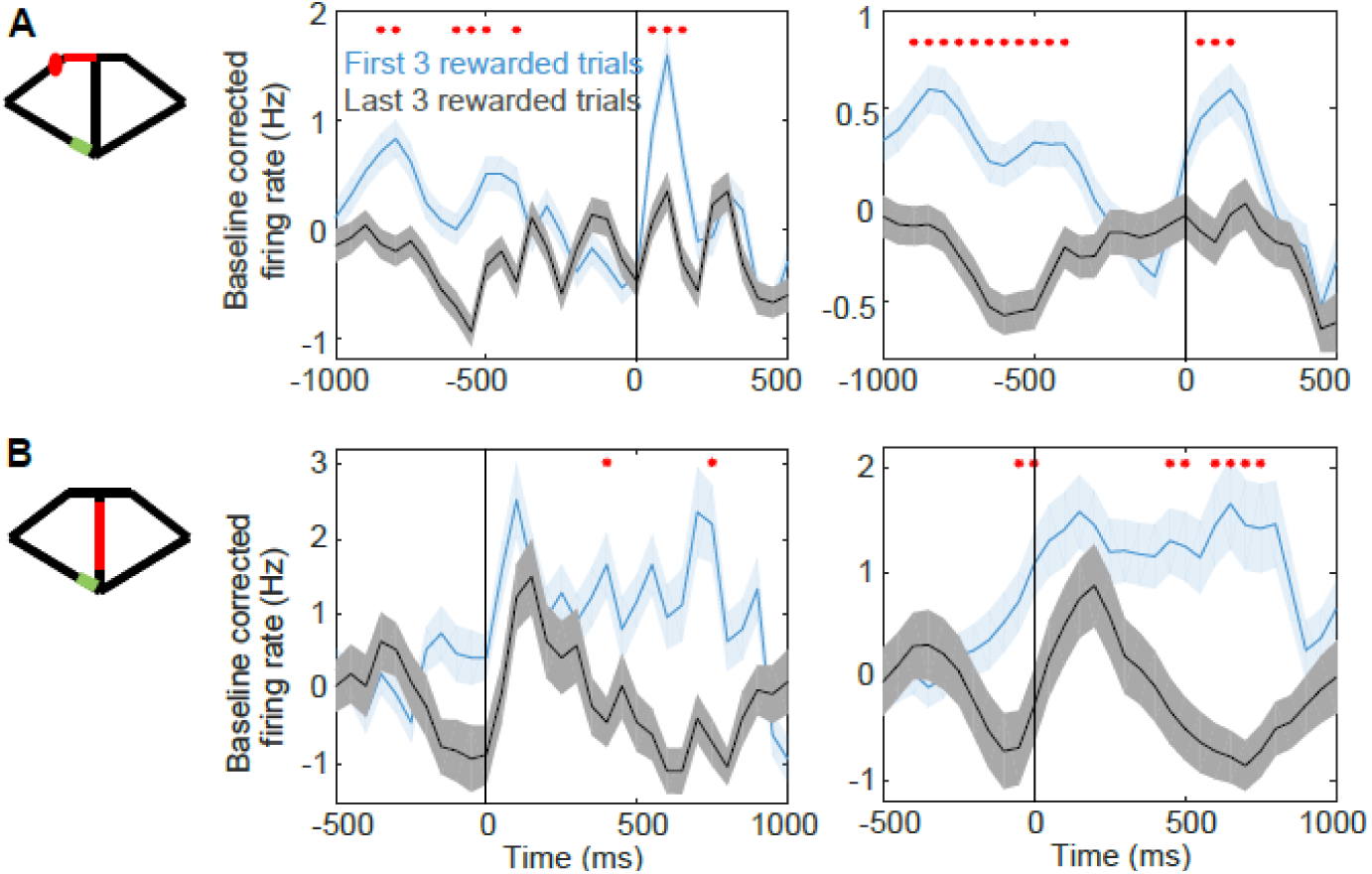
The dopamine response to the earlier cue becomes more phasic after extended training. (A) Dopamine response to the reward onset at the beginning and at the end of each block (4 rats, 62 dopamine cells). The inset shows the experimental apparatus. Firing rates in the red region of the inset were normalized by subtracting the mean firing rate in the green region. The reward was delivered in the region marked with a red circle in the inset. (B) Dopamine response to the bridge lowering. First 3 rewarded (light blue) and last 3 rewarded (gray) trials of the experiment are shown. The right plots show smoothed dopamine response on the left plots. A moving average window of 5 was used. Red asterisks indicate significant differences between the light blue and gray graphs (paired t-test; p < 0.05). Shading shows mean ± SEM.

Since Collins et al. [34] used a lever-press experiment while Howe et al. [46] utilized a maze paradigm, it is uncertain if extended training in a maze paradigm would transform dopamine activity from ramping into phasic. To test this possibility, we compared predictions of our model with unpublished data collected from a maze task (see Acknowledgement). In this experiment, rats freely chose between two arms of a modified T-maze (Figure 4AB inset). The reward probability of one arm was higher than that of the other, and these probability values remained constant within a block of 17-72 trials (mean ± SD: 44.1 ± 12.0). The arm-reward relationship was reversed across four blocks without any sensory cues indicating this change. To control the inter-trial interval, a connecting bridge was lowered 2 s after the animal arrived at the entrance of the bridge (red in Figure 4B inset). Therefore, the lowering of the bridge functioned as a trial initiation cue. Note that although this study recorded electrophysiological activity of dopamine neurons while Howe et al. [46] and Collins et al. [34] used fast-scan cyclic voltammetry (FSCV) in the ventral striatum, previous studies have shown that dopamine concentration kinetics in the ventral striatum is similar to the kinetics of electrophysiological dopamine activity [10,49–51].

As learning proceeded, the dopamine response to the reward onset diminished (Figure 4A). Specifically, as the dopamine activity between the bridge lowering and the reward onset decreased, the dopamine response to the bridge lowering became more phasic (Figure 4AB). These findings are consistent with the prediction of our model that the prediction error gradually backpropagates to the earlier cues and the shape of the prediction error transitions from ramping to phasic during RL. While the dopamine excitation at the reward onset decreased towards the end of each block, about four blocks were needed for the dopamine activity after the bridge lowering to significantly decrease. This is consistent with the results of Collins et al. [34], a recent experimental finding [52], and our simulation that a large phasic response to the initial cue necessitates extended training. Therefore, our model clearly replicates key features of the dopamine activity transitions from ramping to phasic.

### The transition from phasic to ramping patterns upon changes in reward value

According to our model simulation (Figure 3AB), prediction errors around the reward onset diminish as learning proceeds; this indicates a reduced sensitivity to the change in the reward value. Figure 5A shows that the prediction error at the time of reward delivery is larger when the size of the reward is doubled during early training (when the prediction error signal exhibits a ramping pattern) than late training (when the prediction error signal exhibits a phasic pattern). This simulation result indicates that extended training transforms dopamine activity from a ramping pattern to a phasic excitation to the initial cue. Further, it suggests that phasic dopamine focuses cognitive resources on the initial cue at the expense of reduced sensitivity to environmental changes [53]. It is consistent with previous findings that extended training makes learned responses habitual and less flexible [54,55]. Recent studies have found that dopamine activity is closely linked to initiating a movement or a sequence of learned actions [8,9,19,23,24,52,56–58]. These findings, along with our simulation, suggest that during early training when an animal is uncertain about the key stimuli for the given task, many effective task dimensions—each generating a weak prediction error signal and affecting animal’s behavior for a short while—guide the animal to complete the task. However, after extended training converts dopamine activity into a phasic pattern, the initial cue automatically triggers the whole behavior sequence required to complete the task [34,59].

**Figure 5.**
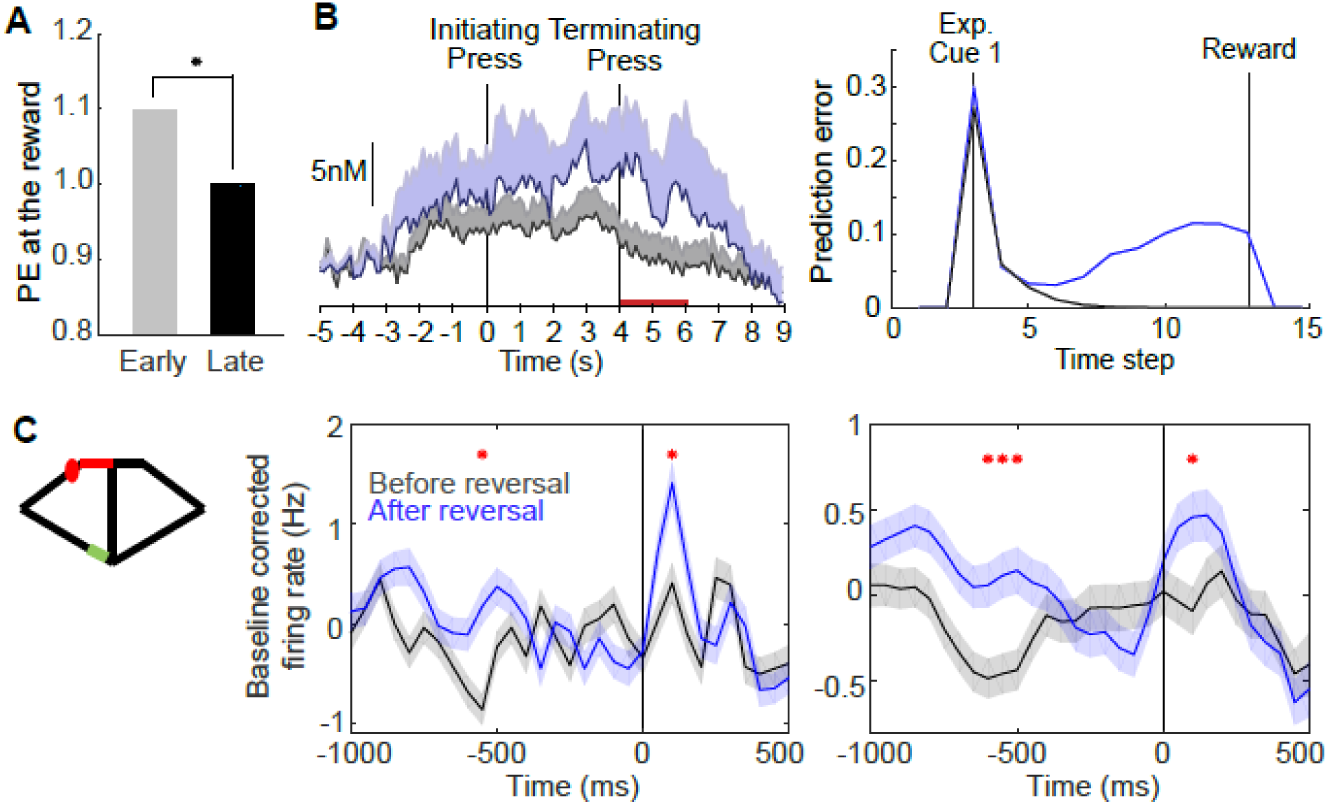
Dopamine transitions from phasic to ramping upon value changes. (A) Prediction error at reward delivery when the size of the reward was doubled during early or late training. We ran 100 simulations with different initial weights (random weights with a mean of 0.02). In each simulation, the first prediction error after a reward increase was calculated. The black asterisk indicates a significant difference (unpaired t-test; p ≈ 0) (B). The experimental results of Collins et al. [34] (left) and the model behavior (right). After extended training, the ramping activity disappeared (gray). The ramping activity reappeared when the size of the reward was doubled (blue) after extended training in Collins et al. [34] (left) or during late training in the model (right). The left plot was originally published in Collins et al. (2016) [34] under a CC-BY 4.0 license. It’s adapted here with permission. The right plot shows prediction error during 10th percentile of training after the reward increase. We ran 100 simulations with different initial weights. Shading shows mean ± SD. (C) Dopamine activity before and after reversal (4 rats, 62 dopamine cells). Red asterisks indicate significant differences between the blue and black graphs. (paired t-test; p < 0.05). Shading shows mean ± SEM. All data are available in the Figure 5_source data 1.

Since phasic activity locked to a particular stimulus is not suitable for fast adaptation to environmental changes, we predicted that, when there is a change in the reward structure, the ramping pattern would reappear, facilitating re-identification of reward-predicting stimuli. Both previous experimental results and our simulation results support this prediction. In particular, our simulation (Figure 5B right) replicated the findings of Collins et al. [34] that ramping activity reappeared when the reward value was doubled after extended training (Figure 5B left; [34]). A similar effect was found in the unpublished data (Figure 5C). After the reward probabilities of the left and right arms were reversed, dopamine activity before the reward onset increased. Overall, our results show that transitions from phasic to ramping activity reflect widespread resource distribution, which allow rapid adaptation to potential changes in the stimulus value. Recent studies have found that the dopamine neurons are responsive to novel stimuli [60,61], changes to reward features [62,63], and stimuli that are potentially relevant to a given task [2–4,64–66]. These findings, together with our simulation, suggest that dopamine activity is suited to adjusting effective task dimensions in response to environmental changes.

## Discussion

In the present study, we tested the theoretical possibility that dopamine transitions between ramping and phasic patterns during RL reflect efficient resource allocation. Both the simulation and experimental results support the view that dopamine activity transitions from ramping to phasic as the RL agent narrows down the candidate reward-predicting stimuli to decrease the effective task dimensionality. The opposite occurred when the agent had to re-identify task-relevant stimuli by increasing the effective task dimensionality. This affords insight into a more fundamental question: how do animals resolve the tradeoff between prediction performance and efficient resource management?

Our model provides a testable prediction that the magnitude of the prediction error protrusion at the intermediate cue depends on the saliency contrast between experimental and non-experimental cues (Figure 6A). Our model also predicts that, the more salient a task-irrelevant stimulus, the more likely a learning agent would mistakenly associate the stimulus with a reward. Previous studies have suggested that the saliency of a stimulus is influenced by multiple factors including the stimulus’ value, a sudden change in the value, uncertainty, or sensory intensity [5,33,67–71]. Since we successfully accommodated the effect of saliency by making parsimonious modifications to a standard TD model, our theory warrants the replication of a wide range of experimental data.

**Figure 6.**
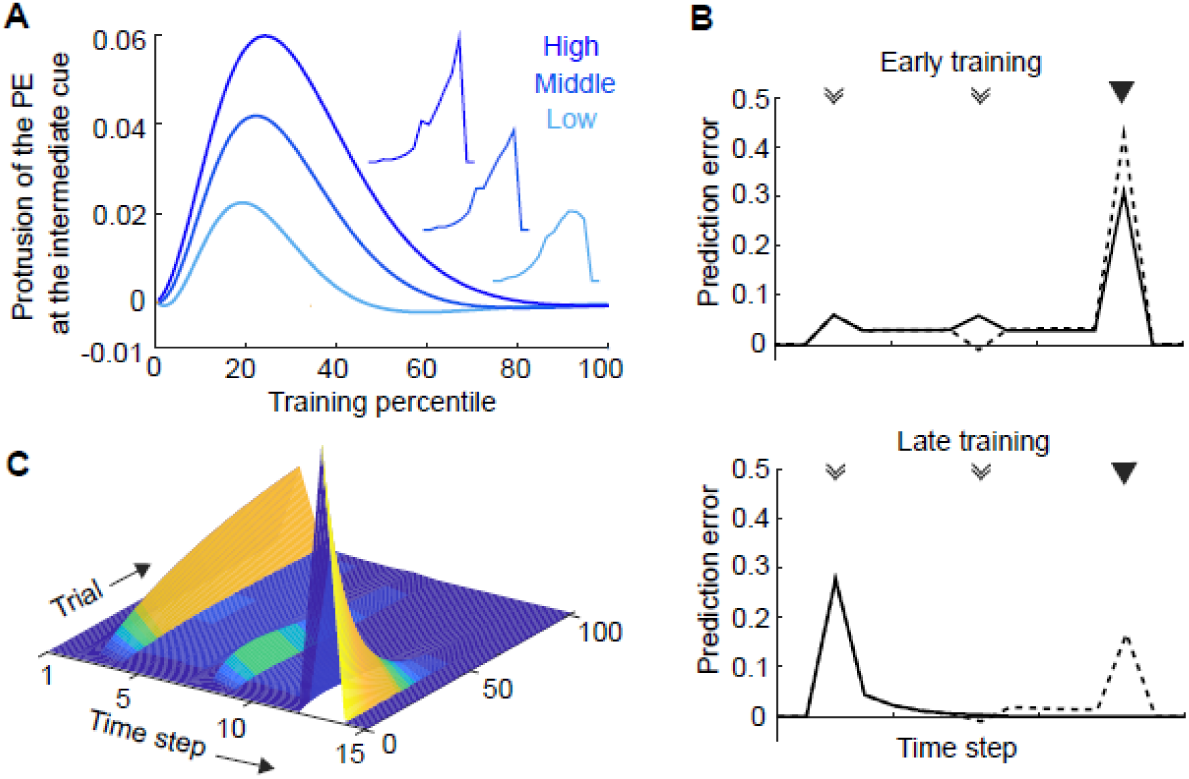
Model prediction and application to simple conditioning experiments. (A) The model predicted that as the saliency contrast between experimental and non-experimental cues increase, the prediction error at the intermediate experimental cue would protrude accordingly. The saliency of the experimental cues was 2, 4, and 8 for high, middle, and low contrast, respectively, while the saliency of the non-experimental (environmental) cues was 1. The level of the protrusion of the prediction error at the intermediate experimental cue was measured as the prediction error at the intermediate experimental cue minus the average of the prediction error immediately before and after the experimental cue. The inset shows the prediction error trajectory during early training. λ of 0.5 was used. (B) The simulation results of the model during early and late training. (C) The prediction error signal throughout a simple conditioning experiment with two experimental cues. λ was 0.9 in (B) and (C).

Moreover, our model can also be applied to a simple classical conditioning experiment, such as Pan et al. [36]. In that experiment, rats were placed in a small, simple chamber and two consecutive tone cues deterministically predicted a liquid reward. Animals only had to lick a spout to obtain the reward. This situation corresponds to the case with a large λ because effective task dimensionality is likely to be smaller than the working memory capacity (refer to Section *Limited resources and environmental stimuli…*). During early training, dopamine neurons showed strong phasic excitation to the reward, whereas the strong dopamine excitation was transferred to the initial cue during late training. Regardless of the learning stage, the omission of the second experimental cue resulted in a larger phasic response to the reward (Figure 6B of [36]). All the dopamine activity patterns were successfully simulated by our model (Figure 6BC).

We chose the value of λ based on the presumed ratio between effective task dimensionality and the working memory capacity. According to this logic, λ should increase as learning proceeds because effective task dimensionality decreases during RL. However, we did not implement this effect and fixed the value of λ throughout each simulation for model simplicity. Adding this effect would have accelerated learning but would not have changed the major findings of the present study.

Recent empirical studies have found an increasingly diverse repertoire of dopamine activity that cannot be interpreted as a prediction error signal [17,58–63,67–70]. Our hypothesis examining the role of dopamine in resource allocation provides a framework to understand the diverse dopamine activities; prediction error is a major (but not the only) factor that determines resource allocation.

Overall, the present study provides a potential explanation for how resource allocating dopamine deals with the task dimensionality problem that RL has been known to be notoriously inefficient at tackling. Further works should investigate more fundamental problems, including the role of dopamine in resolving the tradeoff between reward maximization and resource consumption minimization.

## Materials and Methods

To investigate if dopamine adjusts effective task dimensionality during RL, we incorporated environmental stimuli into a standard TD model with an eligibility trace (Figure 1C). The TD model with an eligibility trace has been shown to well account for dopamine activity during RL [36,37,52]. The goal of a TD learning agent is to maximize the expected amount of future reward. Stimulus value estimation is performed by minimizing the prediction error δ:

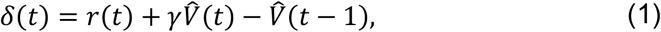

where r(t) is the reward delivered at time *t* and γ is a discounting factor (0 < γ < 1) that decreases the value of delayed rewards. Here, γ was 0.9 in all simulations, and negative prediction errors were bounded by −0.01 [36]. V(t) is a value function that represents the expected value of the temporally discounted sum of all future rewards:

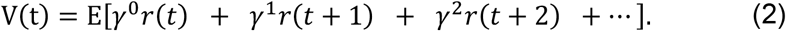

Each stimulus k contributes to the estimate of *V(t)*, and the future reward estimated from each stimulus k is an inner product of the respective state vector x_k_(t) and the weight vector w_k_(t):

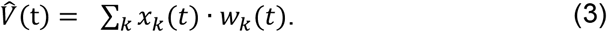

If stimulus k occurs at time n, (n + m)-th element of x_k_(n + m) is 1 (where *m* = 0, 1, 2, 3…) and all other elements of x_k_ are 0. The state vector x_k_ enables the stimulus *k* that occurs at time n to influence the estimate of *V(n + m)*. The weights are updated as follows:

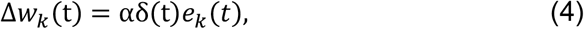

where e_k_ is the eligibility trace for stimulus k [36] and α is the learning rate (0 < α ≤ 1). In all simulations, the learning rate α was fixed at 0.005. The eligibility trace is associated with working memory capacity [43–45]. It quantifies the degree of influence of a prediction error at time t on the value updates in previous time steps and is defined as follows:

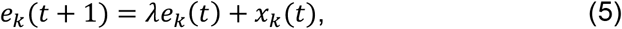

where λ is the eligibility trace parameter (0 ≤ λ ≤ 1). λ = 1 indicates an infinite working memory capacity; the current prediction error updates the values of all previous stimuli. A small λ indicates a small working memory capacity. Thus, using a small λ accommodates a situation in which effective task dimensionality is relatively larger than the agent’s memory capacity (i.e. when many stimuli are being considered for the task). To prevent the prediction errors from carrying over to subsequent trials, the eligibility trace was reset for each trial.

Although experimentally inserted cues (e.g. tones, light) are usually more salient than others (e.g. wells, floor), pseudo-conditioning or generalization indicates that animals also associate the latter with the reward [2–4]. Previous studies have suggested that the more salient a stimulus, the more readily it should be learned [5,69,75]. In the Pearce-Hall model, the constant representing the intrinsic saliency (e.g. intensity) of the cue controls the learning rate [5,68]. To accommodate this in the current study, a saliency signal was incorporated into Equation (4) as follows:

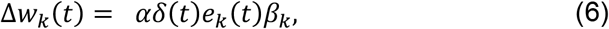

where β_k_ denotes the saliency of the cues. In all simulations, β of experimental and non-experimental stimuli were 2 and 1, respectively. The agent was assumed to complete training when the value of the first experimental cue converged. To clearly demonstrate how the shape of the prediction error signal changed as learning proceeded, the 20th percentile of training—when the prediction error peaked at the second cue—was defined as the middle stage and the last 20th percentile was used to simulate the late stage of learning. Accordingly, the halfth of the middle stage, the first 10th percentile of training, was used to simulate the early stage of learning.

## Acknowledgment

We thank Dr. Min Whan Jung and Dr. Sue-Hee Huh for their generosity in allowing us to use their unpublished data for this paper. We also thank Jung Hwan Shin for pre-processing the unpublished data.

## Competing interests

The authors declared that no competing interests exist.

MRS contributed to conceptualization, methodology, software, investigation, formal analysis, visualization, and writing.
SWL contributed to conceptualization, funding acquisition, project administration, resources, supervision, and writing.

